# Development of functional topography of the default mode subnetworks revealed by precision mapping

**DOI:** 10.64898/2026.02.24.707618

**Authors:** Ying He, Jonah Kember, Hongxiu Jiang, Xiaoqian J. Chai

## Abstract

The default mode network (DMN) is a functionally and anatomically heterogeneous network comprising distinct subsystems that support internally directed processes such as memory and self-related thought. Specialization and segregation of functional brain networks are critical for cognitive maturation, yet how DMN subnetworks develop remains unclear. Group-average approaches obscure individual variability and blur spatial details, limiting precise characterization of network topography. Using precision mapping in 547 participants aged 5–21 years, we delineated individualized DMN subnetworks and examined their development across multiple aspects, including functional segregation and spatial topography. We found that DMN subnetworks become increasingly functionally segregated from each other and from non-DMN networks with age, and exhibit greater topographic distinctiveness through boundary sharpening and reduced spatial overlap. Furthermore, we observed a selective spatial contraction of the memory-related subnetwork, which correlated with better episodic memory. Our findings underscore the importance of characterizing multidimensional functional and topographical features at the individual level to better understand typical neurodevelopment and neurodevelopmental and psychiatric conditions.

## Introduction

The default mode network (DMN) comprises interconnected and distributed brain regions and is thought to be involved in internal mental processes, such as remembering, thinking about future, and self/other-related thoughts^1, 2^. Disruptions of the DMN have been shown in many neuropsychiatric and developmental disorders^3, 4, 5^. Emerging evidence suggests that the DMN is a complex network that consists of distinct subnetworks, each supporting specific cognitive processes^6, 7, 8^. During child development, the broader functional brain networks become increasingly segregated—characterized by stronger within-network and weaker between-network connectivity)^9, 10, 11^. The DMN as a whole becomes more internally integrated while becoming increasingly segregated from other functional networks during development^12, 13, 14^. This growing segregation supports cognitive development by enabling more specialized network functioning^13, 15^. However, how functional segregation between distinct DMN subnetworks develop in children remains unknown. Delineating the development of DMN subnetworks in a neurotypical population is essential, as it provides a crucial baseline for understanding alterations in related neuropsychiatric disorders.

To date, investigations of functional networks in children have largely relied on group-average (one-size-fits-all) parcellations. These group-average approaches introduce spatial blurring and fail to capture individual-specific features of brain organization. This limitation has motivated the growing use of precision mapping approaches, which identify functional networks at the individual level. By capturing individual variability, these methods offer improved reliability in linking brain organization to behavior and symptoms^16, 17^. Furthermore, inter-individual variability in functional brain networks may change across development^18^. In addition, studies using within-person precision mapping approaches have shown that the individual topography of functional networks — including their size, shape, and spatial location — differs markedly from the group average^19, 20^. These topographical variations become more refined during youth development and are associated with individual differences in cognition^21, 22^, with comparable alterations reported in clinical populations^23, 24^. Thus, in addition to assessing within- and between-network connectivity strength and other graph-based measures, this approach also enables the examination of individual topographic features and spatial extent of DMN subnetworks.

In the present study, we performed individual-level network parcellation in a cohort of 547 participants aged 5–21 years, each with more than 21 minutes of resting-state fMRI data. Using the Yeo 17-network template as a reference, we identified three distinct DMN subnetworks (DMNA, DMNB, DMNC) based on individual functional connectivity profiles at the vertex level. A previous meta-analysis of the functional profiles of the three DMN subnetworks found that DMNA was more strongly associated with self-related processes, emotion and evaluation, as well as social and mnemonic processes shared with the other two subnetworks^25^. DMNB was linked to mentalizing and social cognition, as well as story comprehension and semantic/conceptual processing. In contrast, DMNC showed strong associations with past and future autobiographical thought, episodic memory, and contextual retrieval. To understand the development of these three DMN subnetworks, we first examined functional segregation of the three DMN subnetworks, quantified by the segregation index derived from the average within-network and between-network functional connectivity. This allowed us to assess whether the subnetworks became increasingly segregated from one another and from other non-DMN networks with age. Second, we mapped the topography of the three subnetworks individually and examined the topographical segregation, measured by boundary refinement and spatial differentiation between the subnetworks. Third, we examined age-related changes in the surface area of each subnetwork to determine whether their cortical representations expanded or contracted during development. After examining age effects on these segregation indices and spatial extent, we tested whether these developmental changes in brain network organization could predict corresponding cognitive development. Given that DMNC is relatively specific to episodic memory, we examined correlations between memory performance and DMNC metrics.

## Results

### DMN subnetworks show increasing functional segregation with age

Here we employed a template-matching method to obtain individual parcellations of three DMN subnetworks (Figure 1). After obtaining the individually mapped functional networks, we first calculated the within-network homogeneity and compared it to that obtained using a group-level template. The homogeneity of individual networks was significantly higher than that of the group template (*t*_546_ = 15.42, *p* < 0.001), indicating the effectiveness of our precision mapping method. In addition, we computed the Adjusted Rand Index between each subject’s parcellation under a given threshold and the parcellations of the same and other subjects under different thresholds. We observed that parcellations from different thresholds within the same subject showed higher similarity than those between different subjects (Figure S1), suggesting that our precision mapping method produces consistent results within individuals and is robust to parameter variations.

**Figure 1.**
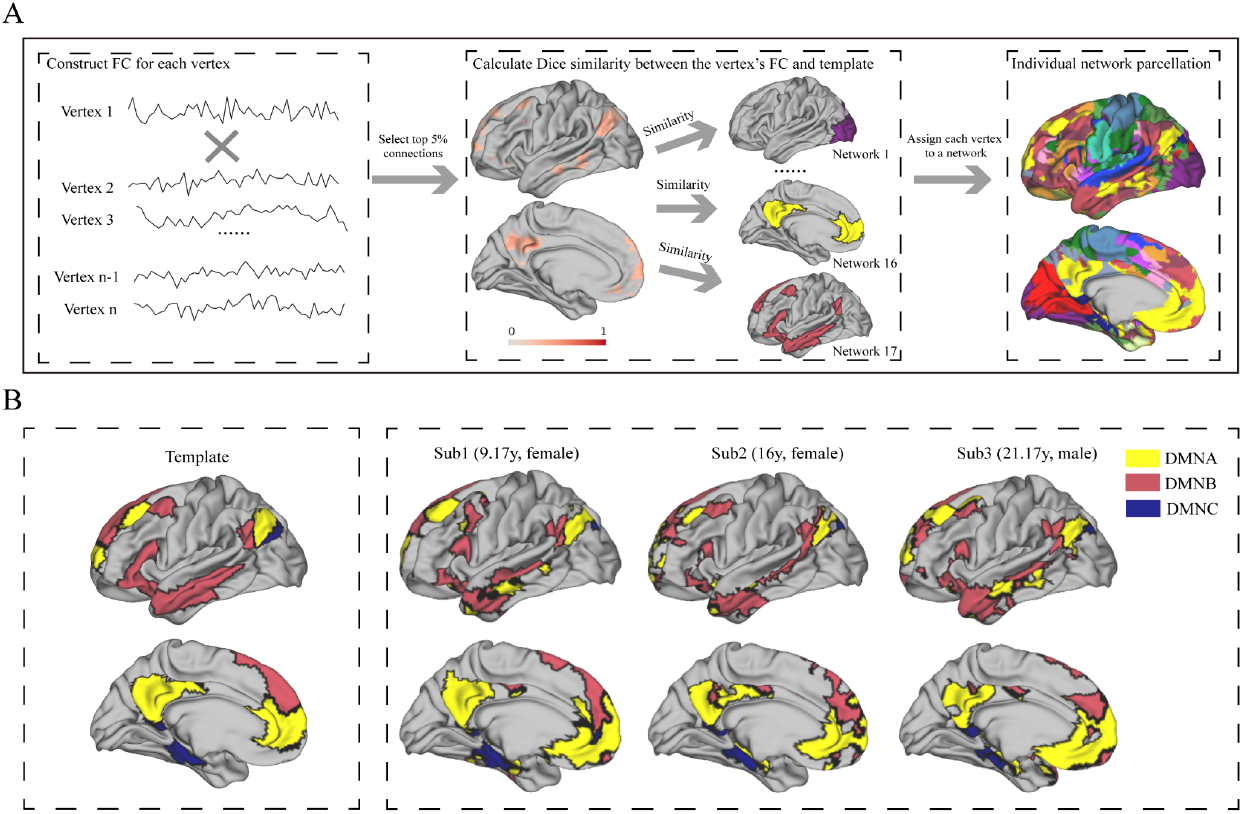
Overview of the precision mapping method. A. First, we constructed the functional connectivity profile for each vertex by calculating the Pearson correlation between that vertex and all other vertices. Next, we computed the Dice similarity metric to quantify the spatial similarity between each vertex’s connectivity profile and each of Yeo’s 17 networks. Finally, each vertex was assigned to the network with which it showed the highest similarity. B. The left panel shows the template of the DMN subnetworks based on the Yeo 17-network parcellation. The right panel displays three representative participants selected from different age groups, illustrating individual differences in the topographical distribution of the DMN subnetworks.

We quantified functional segregation by computing a segregation index based on within-network and between-network functional connectivity. Within the DMN, each subnetwork showed significantly increased functional segregation from the other two subnetworks with age (Figure 2A, DMNA: *β* = 0.005, *t*_541_ = 6.37, *p*_corrected_ < 0.001; DMNB: *β* = 0.008, *t*_541_ = 7.65, *p*_corrected_ < 0.001; DMNC: *β* = 0.008, *t*_541_ = 6.76, *p*_corrected_ < 0.001). These findings demonstrate progressive functional segregation among all three DMN subnetworks across development.

**Figure 2.**
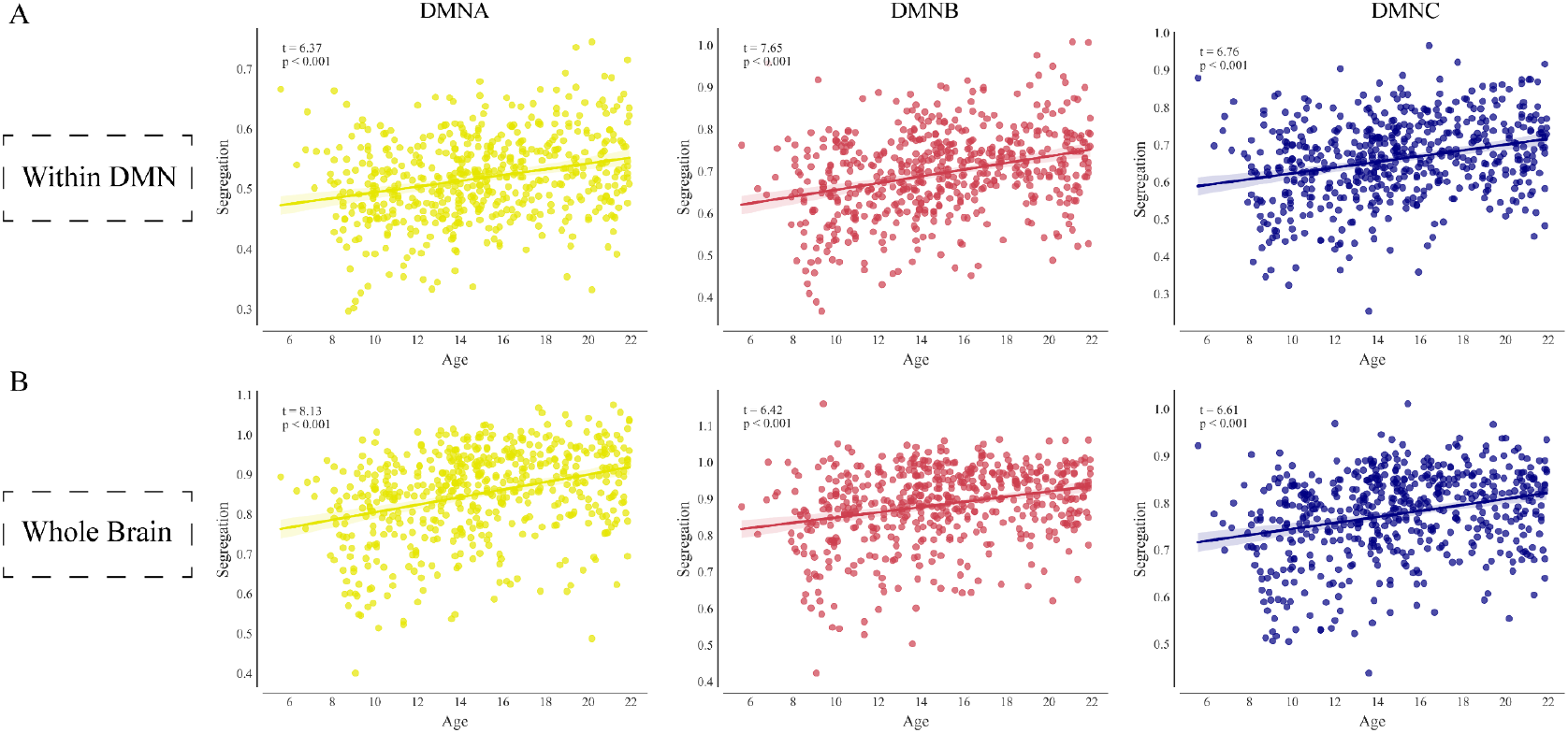
DMN subnetworks show increasing functional segregation across development. A. The segregation of each DMN subnetwork relative to other DMN subnetworks significantly increased with age. B. The segregation of each DMN subnetwork from all other large-scale functional networks also showed a significant age-related increase. Shaded bars reflect 95% confidence intervals of the regression line.

In addition, we examined the age-related changes in functional segregation between each DMN subnetwork and the other 16 large-scale functional networks. Similar to segregation relative to other subnetworks of DMN, all DMN subnetworks showed significant increases in segregation from the other 16 networks with age (Figure 2B, DMNA: *β* = 0.010, *t*_541_ = 8.13, *p*_corrected_ < 0.001; DMNB: *β* = 0.007, *t*_541_ = 6.42, *p*_corrected_ < 0.001; DMNC: *β* = 0.006, *t*_541_ = 6.61, *p*_corrected_ < 0.001). We also tested whether increasing functional segregation of DMNC from the other two DMN subnetworks and from all other 16 networks correlated with PSM scores, but no significant correlations were found (all *ps* ≥ 0.42). These findings further highlight the developmental emergence of functional differentiation of DMN subnetworks.

### Topographical Boundaries of DMN Subnetworks Become Sharper with Age

Here, we examined the spatial distribution of the three DMN subnetworks among the participants using the density map (Figure 3A). We found that the highest density regions of each subnetwork were primarily located within their respective canonical cortical zones, but also extended into regions typically associated with other subnetworks, albeit at lower densities. For instance, the posterior cingulate cortex and precuneus regions were typically assigned to DMNA, although some individuals showed assignments to DMNB. The anterior temporal regions were generally associated with DMNB, but a substantial proportion of participants had anterior temporal regions that were assigned to DMNA. Similarly, the ventral medial prefrontal cortex was most often assigned to DMNA, yet assignments to DMNB and DMNC were also observed in some individuals. This pattern reflects both individual variability and their shared characteristics, with the three networks being distributed yet adjacent.

**Figure 3.**
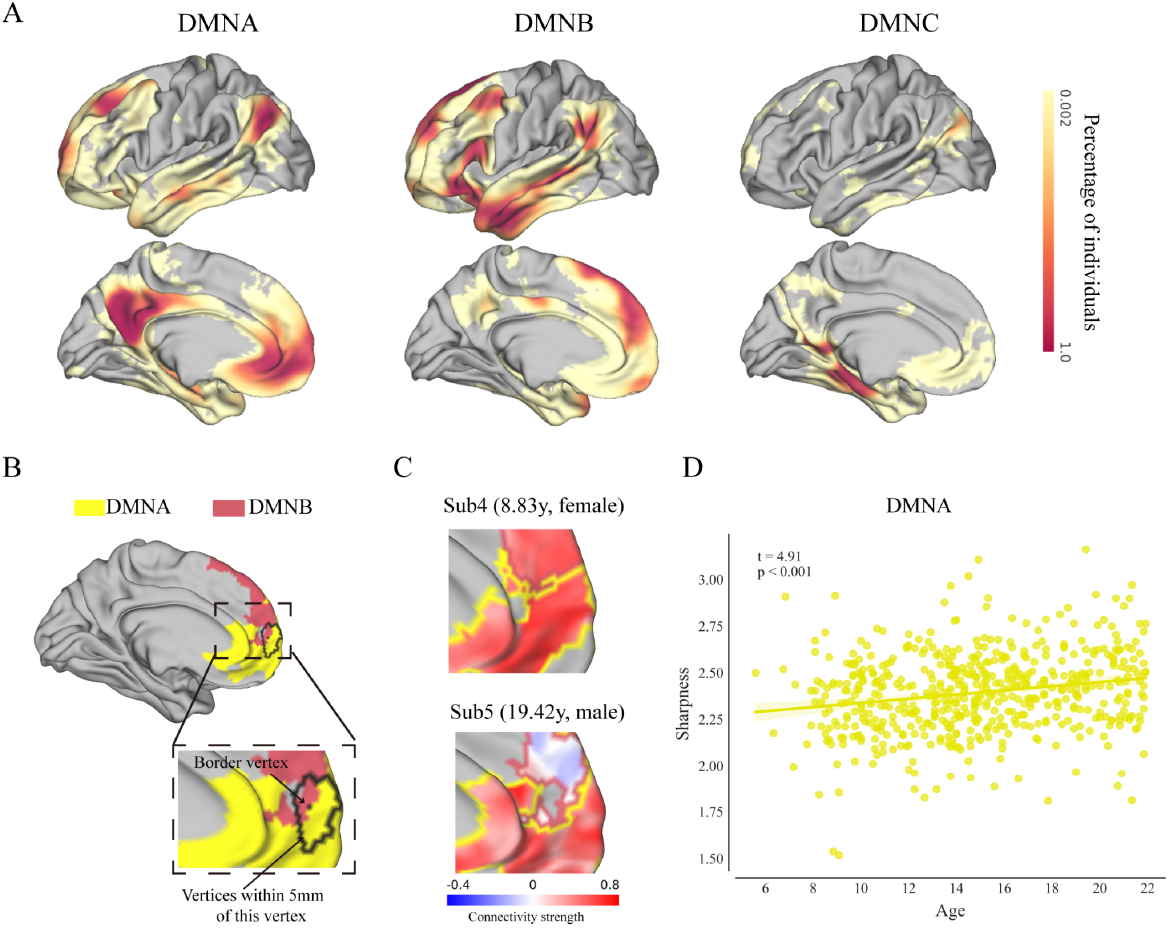
Topographical boundaries of DMN subnetworks became sharper with development. A. The density maps indicated the proportion of subjects for whom a given vertex was assigned to each of the three DMN subnetworks. The spatial locations of the three DMN subnetworks were primarily centered around their respective centroids within each cortical zone, but they also extended into regions typically associated with other subnetworks. B. Two clusters from different DMN subnetworks were selected as examples to illustrate how boundary sharpness was quantified. For each border vertex between the two clusters, we identified all vertices located within 5 mm and used them for the analysis. C. Two example subjects illustrating boundary sharpness between DMNA and DMNB. Red outline indicates the DMNB cluster, and the yellow outline indicates the DMNA cluster. The color bar represents the functional connectivity strength of each vertex to the mean time series of the DMNA cluster. The top panel shows a subject with a blurred boundary, where vertices along the border exhibit highly similar connectivity to both subnetworks. The bottom panel shows a subject with a sharper boundary, where border vertices display distinct connectivity profiles between the two subnetworks. Note that while most cluster vertices are shown for visualization, sharpness was calculated only for border vertices within 5 mm (panel B). D. The scatter plot shows that DMNA boundaries with both other subnetworks became significantly sharper with age.

Furthermore, we aimed to investigate whether the topographical boundaries between DMN subnetworks become sharper with age. To do so, we first identified boundary vertices based on their proximity (within 5 mm) to the borders between adjacent subnetwork clusters (Figure 3B). Boundary sharpness was defined as the extent to which these vertices showed stronger functional connectivity with their own subnetwork compared to the neighboring one, indicating a clearer distinction between adjacent subnetworks. Here, we present two examples illustrating a blurred versus a sharp boundary (Figure 3C). We examined age-related changes in mean boundary sharpness for each DMN subsystem, calculated by averaging the sharpness values of the boundaries between the given subnetwork and the other two subnetworks. DMNA showed a significant increase in mean boundary sharpness with age (*β* = 0.01, *t*_541_ = 4.91, *p*_corrected_ < 0.001, Figure 3D), driven by sharpness increases at both its interfaces with DMNB (*β* = 0.01, *t*_541_ = 3.65, *p*_corrected_ < 0.001) and DMNC (*β* = 0.01, *t*_541_ = 2.78, *p*_corrected_ = 0.010, Figure S2). We didn’t find significant age-related changes in mean boundary sharpness in DMNB (*β* = 0.006, *t*_541_ = 1.77, *p*_corrected_ = 0.114) or DMNC (*β* = 0.002, *t*_541_ = 0.34, *p*_corrected_ = 0.731) after FDR correction. This pattern of results suggests that DMNA boundaries undergo progressive sharpening during development, whereas DMNB and DMNC boundaries remain relatively stable.

### Spatial overlap of DMN subnetworks decreases with age

After demonstrating the refinement of boundaries with age, we next sought to investigate the spatial differentiation among DMN subnetworks. We defined spatial overlap between DMN subnetworks as vertices that simultaneously showed high functional connectivity with the mean time series of two subnetworks, where “high” was defined as connectivity exceeding the subject-specific mean by at least one standard deviation. We then examined how the surface area of these overlapping regions changed with age. Among the subnetworks, the spatial overlap between DMNA and DMNB exhibited the highest surface area proportion, and this overlap was significantly negatively correlated with age (*β* = -0.003, *t*_541_ = -2.53, *p*_corrected_ = 0.017, Figure 4A). Similarly, the overlap between DMNB and DMNC also showed a significant negative correlation with age (*β* = -0.002, *t*_541_ = -2.71, *p*_corrected_ = 0.017, Figure 4B). The spatial overlap between DMNA and DMNC did not show a significant relationship with age (*β* = 6.96E-05, *t*_541_ = 0.07, *p*_corrected_ = 0.947). Additional analyses across different correlation thresholds for defining high spatial overlap (from the subject-specific mean, to two standard deviations above) confirmed the robustness of the age-related decrease in overlap between DMNA and DMNB (Figure S3). These findings indicate that the spatial distribution of the DMN subnetworks becomes more differentiated during development, especially between DMNA and DMNB.

**Figure 4.**
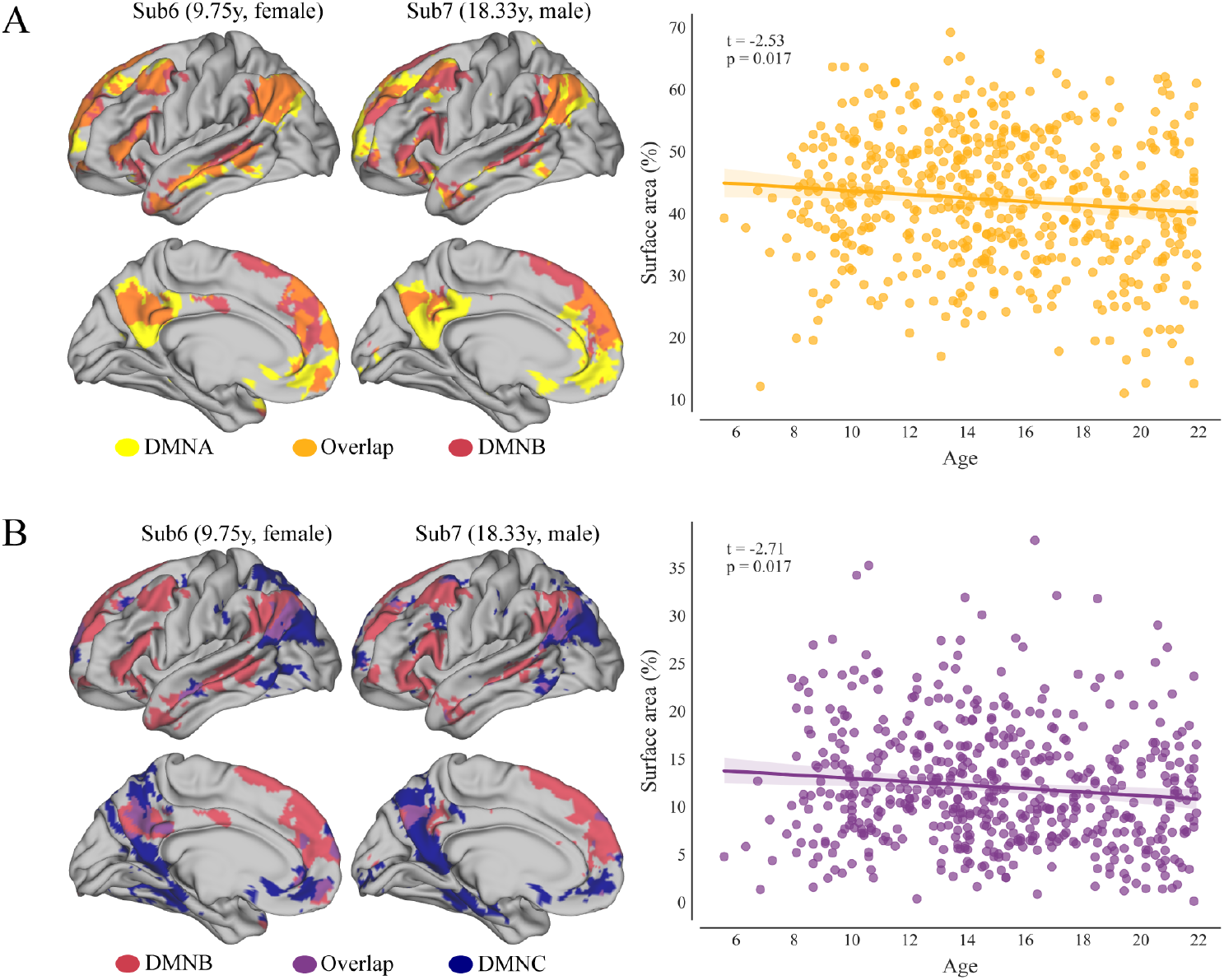
Overlap of DMN subnetworks decreased with age. A. Two examples illustrate the spatial overlap between DMNA and DMNB from a younger and an older participant, respectively. Only the left hemisphere is displayed for visual clarity. Analyses were conducted on both hemispheres. The accompanying scatter plot demonstrates a significant decrease in overlap with age. B. Two examples show the spatial overlap between DMNB and DMNC from a younger and an older participant. The scatter plot again indicates a significant age-related decline in overlap.

### Surface area of DMNC decreases with age

We first examined age-related changes in the surface area of DMN subnetworks using a GLM, controlling for sex and scanning sites. After correction for multiple comparisons, we observed a significant age-related decrease in the surface area of the DMNC subnetwork (*β* = –0.04, *t*_541_ = –4.00, *p*_corrected_ < 0.001). No significant associations with age were found for the DMNA (*β* = 0.01, *t*_541_ = 0.58, *p*_corrected_ = 0.63) or DMNB (*β* = –0.01, *t*_541_ = –0.94, *p*_corrected_ = 0.45) subnetworks (Figure 5A). When treating the DMN as a whole network, its overall surface area does not exhibit significant age-related changes (*β* = -0.02, *t*_541_ = -1.53, *p*_corrected_ = 0.19). To further illustrate age-related changes in DMNC surface area, subjects were divided into three age groups based on age: children (≤12 years, n=147), adolescents (12-18 years, n=262), and adults (≥18 years, n=138). The corresponding density maps show that as age increases, the surface area of the DMNC gradually decreases and becomes more spatially constrained (Figure 5B). We also observed a significant age-related reduction of DMNC in the ventromedial prefrontal cortex (vmPFC), a region typically associated with the DMNA (Figure 5B). This pattern suggests a spatial reorganization of DMN subnetworks during development, with DMNC territory potentially redistributing to other subnetworks in adulthood.

**Figure 5.**
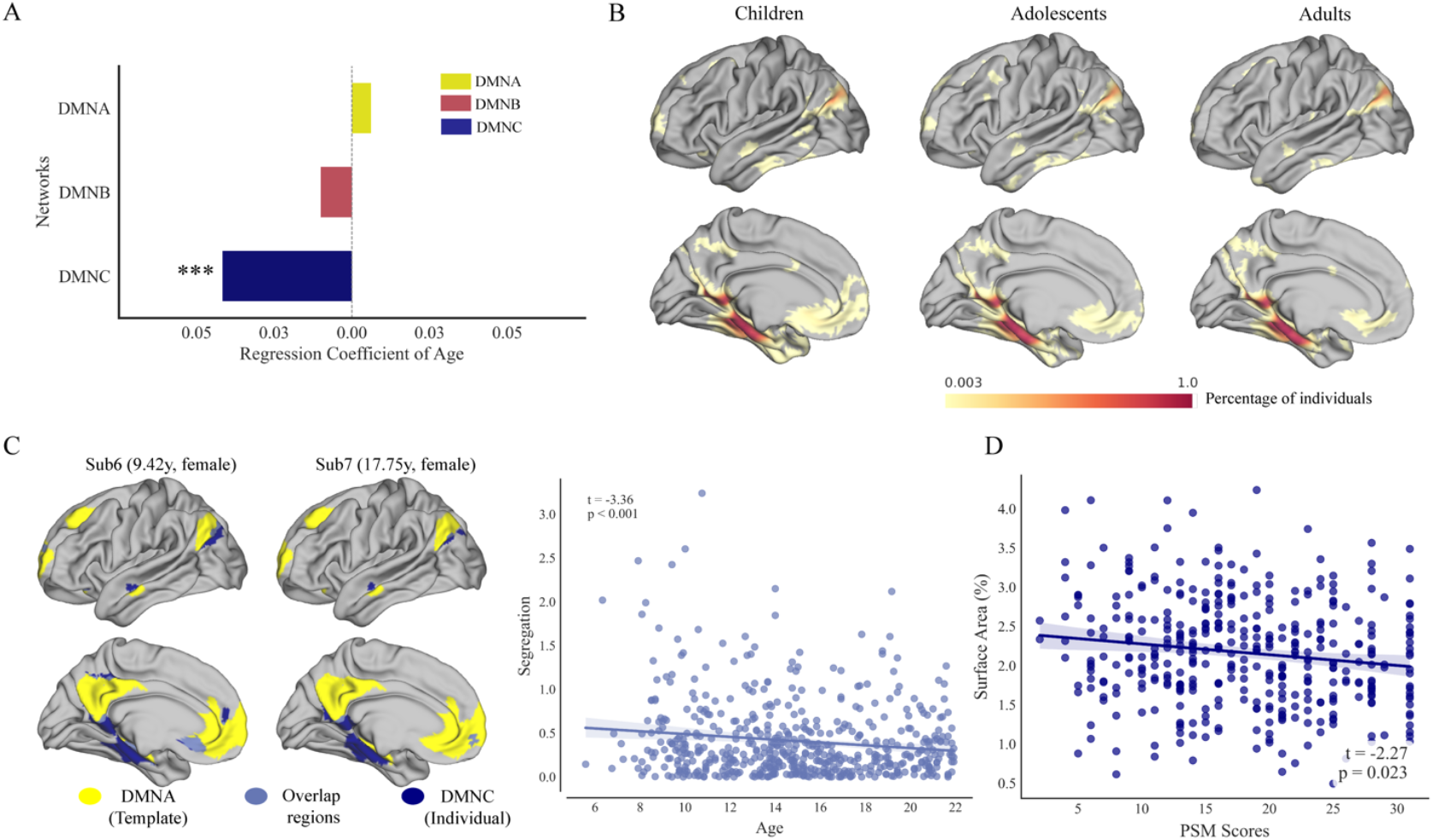
Surface area of DMNC decreased with age and memory performance. A. This bar plot illustrates the effect of age on the surface area of DMN subnetworks, based on a general linear model in which age was treated as the independent variable and surface area as the dependent variable, while controlling for confounding factors (sex, site, etc.). The width of each bar reflects the regression coefficient for age. Asterisks indicate levels of statistical significance after FDR correction: **p < 0.001; *p < 0.01; p < 0.05. B. Panel B shows the density maps for different age groups. Density values represent the proportion of subjects within each group in which a given vertex was assigned to the DMNC. C. The left panel shows two example subjects, illustrating the spatial overlap between individual DMNC (blue) and the group-level DMNA from template (yellow). The scatter plot demonstrates an age-related decrease in the overlap proportion (overlap area / individual’s total DMNC area), indicating that as DMNC shrinks during development, it is primarily the regions that correspond to canonical DMNA that are lost. D. A negative association was observed between the DMNC surface area proportion (individual’s DMNC area / individual’s total cortical area) and Picture Sequence Memory performances.

To further characterize the spatial organization of age-related changes in individualized DMNC, we compared each participant’s DMNC map with the group-level template as a reference. This approach enabled us to identify which networks occupy the regions where the DMNC contracts during development, relative to the canonical network spatial distribution. We first quantified the spatial overlap between each participant’s DMNC and the group-level DMNA. As expected, the proportion of individual DMNC overlapping with group-level DMNA (overlap area / individual DMNC area) significantly decreased with age (β = –0.001, t_403_ = –4.92, p < 0.001, Figure 5C). This indicates that regions initially occupied by DMNC in younger children are progressively reassigned to canonical DMNA territory. In contrast, when examining the overlap between youth DMNC and adult group-level DMNB, nearly all subjects showed minimal to no overlap (Figure S4), confirming that the developmental changes in DMNC are relatively spatially specific to DMNA-associated regions. Collectively, these findings demonstrate age-related spatial reorganization among DMN subnetworks, whereby cortical territories undergo systematic reassignment during development.

To further examine whether the DMNC subnetwork contracting is associated with the development of episodic memory, we also examined the relationship between PSM scores and the surface area of DMN subnetworks, controlling for age, sex, and scanning site. A significant positive association was observed between age and PSM score (*β* = 0.71, *t*_396_ = 8.05, *p* < 0.001), indicating that episodic memory performance improves with development. We found that the surface area of the DMNC subnetwork was significantly negatively correlated with PSM scores (*β* = -1.17, *t*_395_ = -2.27, *p* = 0.02), indicating that a reduced surface area was associated with better episodic memory performance. These findings suggest that during development, DMNC exhibits a more compact cortical organization compared to other subnetworks, which may underlie its link to memory performance (Figure 5D).

## Discussion

In this study, we systematically examined the developmental segregation and specialization of DMN subnetworks in 547 participants aged 5-21 years, using precision mapping to obtain individualized parcellations of DMN subnetworks. Our findings reveal developmental segregation occurring across multiple dimensions. First, the three DMN subnetworks exhibited increased functional segregation from each other and from all other networks with age. Second, topographical segregation was observed: boundaries between DMNA and the other two subnetworks became significantly sharper with age, accompanied by reduced spatial overlap between DMNA and DMNB. Third, we observed a selective contraction of DMNC that correlated with better episodic memory, with the contracted regions predominantly overlapping with canonical DMNA territory. Together, these findings provide a comprehensive characterization of DMN subnetwork development, encompassing progressive functional and topographical segregation, and selective spatial focalization.

How to identify the DMN subnetworks remains an area of active research^26, 27^. A commonly used approach is to divide it into three subnetworks, as proposed by Andrews-Hanna et al. (2010, 2014)^7, 25^, with similar parcellations also reported by Yeo et al. (2011)^28^. DMNA is associated with self-related processes, emotion, and evaluation, while also sharing social and mnemonic functions with DMNB and DMNC. DMNB is linked to social processing, whereas DMNC shows relative specificity to episodic memory^25^. Based on these prior findings, we decided to parcellate the DMN into three subsystems, following Yeo et al. Previous studies have examined the functional segregation of the DMN by treating it as a single network. For example, one study reported increased within-network connectivity of the entire DMN during rest from childhood to adolescence^12^. Similar effects have been observed in children during task states^13^. More recently, a large-scale population study found that functional segregation of the canonical DMN, measured using the same index, increases rapidly with age and peaks at the end of the third decade^14^. Only one previous study has investigated the segregation of DMN subnetworks in a developmental sample, comparing the modularity of DMN subsystems between individuals with autism and neurotypical controls^29^. However, this study employed an ROI-to-ROI approach with relatively limited ROI coverage and the same ROI definitions across all participants, which could have overlooked inter-individual variations and thereby contributed to the finding of no marked age-related changes in the neurotypical group^29^. Our approach overcomes this limitation and reveals developmental effects showing that the DMN, a heterogeneous network as established by functional–anatomical evidence^7, 25, 28, 30, 31, 32^, becomes increasingly heterogeneous and functionally segregated over the course of development. Segregation of associative networks is consistently associated with enhanced cognitive performance, while the loss of such segregation in later life has been implicated in cognitive decline^9, 33, 34, 35, 36, 37, 38^. DMN subnetworks share a common organizational motif that supports internally generated representations, with each network specialized for distinct processing domains^27^, and their segregation with development may reflect the enhancement of specific cognitive functions.

We also observed an age-related increase in segregation in the topography of DMN subnetworks. Our precision mapping results reveal individual parcellations with clear and specific spatial boundaries. Given that the three DMN subnetworks are spatially adjacent to one another we examined the age-related effects on boundary sharpness between subnetworks and found that the boundaries around DMNA become progressively sharper across development. At the same time, we found that the spatial overlap between DMNA and DMNB decreases with age, suggesting that DMNA gradually emerges as a more spatially distinct and clearly delineated subnetwork. Notably, DMNA showed the most pronounced age-related changes in topographical segregation among all three DMN subnetworks. DMNA encompasses the so-called midline core regions of the DMN, including the anterior medial prefrontal cortex, angular gyrus, and posterior cingulate cortex, which are among the most consistently engaged regions within the network^7, 25^. Previous group-averaged studies examining functional connectivity between pairs of DMN nodes have reported age-related increases primarily in midline core regions, which is consistent with our findings^12^. Convergent structural and functional connectivity analyses in prior work also suggest that PCC–mPFC connectivity (largely overlapping with DMNA) represents the most immature link within the DMN in children, further highlighting the pronounced age-related effects^39^. Importantly, previous studies have demonstrated that behavioral phenotypes across cognition, personality, and emotion can be predicted by individual-specific network topography with modest accuracy, comparable to predictions based on connectivity strength^40^. Our study advances this line of research by introducing two novel measures of topographical segregation—boundary sharpness and spatial overlap— that uncover new insights into the developmental segregation of DMN subnetworks. These topographical measures, enabled by within-person precision mapping, may provide new biomarkers for predicting behavioral outcomes and developmental trajectories at the individual level, complementing traditional connectivity-based approaches.

Finally, we examined age-related effects on the surface area of DMN subnetworks. While many studies have investigated developmental trajectories of total or regional cortical surface area based on group averages^41, 42, 43^, research focusing on the surface area of individual functional networks remains limited. One study on human lifespan functional connectome changes, using personalized mapping, reported a slight expansion of the whole DMN size during the first month of life but no significant change thereafter^14^. These findings are consistent with ours. Specifically, when we first tested the proportion of the whole DMN surface area relative to the total cortical surface area, we found no significant age-related effects. This result highlights that treating the DMN as a single entity may obscure the heterogeneous developmental trajectories of its subnetworks. When we examined each subnetwork separately, we found that the surface area of DMNC progressively decreased with age. This reduction was predominantly localized to regions that were more frequently assigned to canonical DMNA, indicating spatial reorganization within DMN subnetworks during development. Critically, this contraction was correlated with episodic memory performance, suggesting that its areal specialization supports ongoing cognitive development.

Our study has several limitations. First, we divided the DMN into three subnetworks; however, the exact number of DMN subnetworks remains unclear due to the complexity of DMN function, and our study does not address this question, which warrants further investigation. Second, when examining DMN-related cognitive performance—for example, the close association between the DMNA and self-related thoughts—we lacked corresponding behavioral measures to explore these relationships, which may require a broader behavioral phenotype. Finally, our discussion of cognitive functions associated with DMN subnetworks primarily relies on previous literature rather than functional validation using specific tasks, which should be addressed in future studies.

In summary, our study establishes a novel, multi-dimensional framework for investigating DMN subsystem segregation during development by incorporating measures of functional segregation, topographical segregation, and spatial extent alterations through individualized precision mapping. We demonstrate that progressive functional segregation of DMN subnetworks is accompanied by boundary sharpening and spatial differentiation—manifested as reduced overlap between subnetworks—as well as selective spatial refinement of DMNC that relates to episodic memory performance. By transcending group-averaged approaches and examining individual DMN subnetworks, our findings emphasize the importance of characterizing multi-dimensional functional and topographical features at the individual level to elucidate network segregation and specialization processes essential for typical neurodevelopment and relevant to neuropsychiatric and developmental disorders.

## Materials and Methods

### Data

All participants are from the Human Connectome Project in Development (HCP-D). HCP-D is a cross-sectional project including participants aged 5–21 years. HCP-D MRI data were acquired at four sites: Harvard University, University of California– Los Angeles, University of Minnesota, and Washington University in St. Louis^44^. Our analysis is based on Release 2.0 (N = 625). All participants provided informed assent or consent. Individuals exhibiting excessive head motion (frame-wise displacement > 0.25 mm for more than one-third of the scanning duration) were excluded from further analysis. The final sample consisted of 547 subjects aged 5 to 21 years (mean age = 15 ± 3.8 years, 285 females). The sample was predominantly White (63.4%), followed by individuals identifying as more than one race (15.8%), Black or African American (10.8%), Asian (7.5%), and Other (2.5%). The median annual household income of the sample was $119,000.

### Image acquisition

MRI scans were acquired at 4 sites on 3T Siemens Prisma scanners with a 32-channel Prisma head coil. T1w images were acquired with a 3D multi-echo MPRAGE (Magnetization Prepared Rapid Gradient Echo) sequence with an in-plane acceleration factor of 2. The scanning parameters included: repetition time of 2,500 ms, echo times of 1.8, 3.6, 5.4, and 7.2 ms, inversion time of 1,000 ms, flip angle of 8 degrees, 208 slices, and a voxel resolution of 0.8 mm isotropic. The fMRI scans are acquired with a 2D multiband gradient-recalled echo (GRE) echo-planar imaging (EPI) sequence (MB8, TR/TE = 800/37 ms, flip angle = 52) and 2.0 mm isotropic voxels covering the whole brain (72 oblique-axial slices). To mitigate bias associated with a specific phase-encoding direction, functional scans were acquired in pairs, with one run using anterior-to-posterior (AP) and the other using posterior-to-anterior (PA) phase encoding. Participants aged 8 years and older underwent 26 minutes of resting-state scanning, consisting of four runs of 6.5 minutes each. For younger participants aged 5 to 7 years, a shorter protocol was used, comprising six runs of 3.5 minutes each, totaling 21 minutes.

### Image preprocessing

In our study, we utilized preprocessed T1-weighted and fMRI images obtained from the NIMH Data Archive (NDA). The data were processed using the HCP-recommended preprocessing pipeline, *fMRIVolume*, which includes realignment, correction for spatial distortions, and registration of the functional images to the structural images with minimal smoothing (only due to interpolation, with no explicit smoothing). Artifact removal was performed using *sICA+FIX*, which regresses out motion- and artifact-related ICA components. Finally, *fMRISurface* was used to project gray matter voxels onto the cortical ribbon and register them to the fs_LR 32k surface space.

### Precision mapping

We employed a template-matching approach similar to that used in previous studies^45^. Compared with other precision mapping methods—such as Infomap, non-negative matrix factorization, and Overlapping MultiNetwork Imaging mapping— template matching requires less scan time to obtain reliable personalized maps^45^. In this study, we used Yeo’s 17-network parcellation as the template. For each vertex, we computed its functional connectivity profile with all the other vertices and retained the top 5% of connections. We also examined the top 1% and 3% of connections and compared the consistency of parcellation across different thresholds using the Adjusted Rand Index, to ensure that the results were not driven by the specific choice of threshold. We then calculated the Dice similarity coefficient between each vertex’s connectivity profile and each of the 17 Yeo networks, generating a vertex-by-network similarity matrix. Each vertex was assigned to the network with which it showed the highest similarity. Subsequently, we applied the Connectome Workbench (https://www.humanconnectome.org/-software/connectome-workbench) command to identify and remove small isolated clusters smaller than 40 mm^2^. These small clusters were then spatially reassigned to the nearest neighboring network to ensure spatial contiguity. Finally, we obtained the individual network parcellation for each subject.

### Homogeneity of functional networks

To evaluate the effectiveness of our precision mapping approach, we calculated the within-network homogeneity, following procedures used in previous studies^40, 46^. Specifically, we computed the Pearson correlation coefficients among all vertices within each network and then averaged these values across all networks for both the precision-mapped parcellation and the group-level Yeo’s 17-network parcellation. A paired t-test was conducted to assess whether within-network homogeneity was significantly higher for individual-specific parcellations compared to the group-level parcellation.

### Functional segregation of DMN subnetworks

To investigate the functional organization of DMN subnetworks, we calculated the functional segregation index, a metric that has been used in previous studies^34^. We first extracted the time series of all vertices and computed pairwise Pearson correlations, resulting in a vertex-wise functional connectivity matrix. Next, we applied Fisher’s z-transformation to the connectivity values. For each network, we calculated the segregation index as the difference between its average within-network connectivity (FC_W_) and its average between-network connectivity (FC_B_), normalized by the within-network connectivity, as shown in the formula. We computed the within-DMN segregation (each DMN subnetwork vs. the other two DMN subnetworks) and whole-brain segregation (each DMN subnetwork vs. all 16 other networks).

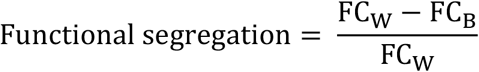

### Density map of DMN subnetworks

To examine individual differences in the topography of DMN subnetworks, we computed the density map of the DMN subnetworks. The density map was generated by calculating the proportion of individuals for whom a given vertex belonged to a specific functional network. Higher values indicate that a greater proportion of participants assigned that vertex to the given network. To better visualize age-related changes in the spatial distribution of functional networks, subjects were divided into three age groups based on age: children (≤12 years, n=147), adolescents (12-18 years, n=262), and adults (≥18 years, n=138). For each group, we generated density maps to visualize the spatial location of the network.

### Sharpness of the boundary between DMN subnetworks

To quantify the topographical distinctiveness of DMN subnetworks, we calculated the boundary sharpness between adjacent networks. First, we used the Connectome Workbench command to identify spatially contiguous clusters within each functional network. Then, we generated borders outlining each cluster, and extracted the vertex indices corresponding to these borders. For each cluster, we computed its mean time series and identified nearby vertices within a 5 mm distance of each border vertex. These nearby vertices were classified as either inside vertices (belonging to the same cluster) or outside vertices (belonging to different clusters). We then calculated the Pearson correlation between the cluster’s mean time series and the time series of each nearby border vertex. To assess boundary sharpness, we computed Cohen’s d for the difference in correlation values between inside and outside vertices. The outside vertices were further categorized into different functional networks based on individual-level precision mapping. This allowed us to compute boundary sharpness between each cluster and each adjacent functional network. If the number of outside vertices belonging to a given network was fewer than 5, the sharpness was not calculated for that network pair. Finally, we averaged the sharpness values of all clusters belonging to the same network to obtain network- to-network sharpness estimates. We separately computed the sharpness between each DMN subnetwork and the other two DMN subnetworks.

### Spatial overlap between DMN subnetworks

To quantify spatial overlap between DMN subnetworks, we built upon the individualized precision mapping results. While precision mapping assigns each vertex exclusively to a single subnetwork through a winner-take-all procedure, functional overlap between subnetworks may exist in practice. To capture this potential overlap, we adopted the following approach. For each subnetwork, we first computed the mean BOLD time series by averaging all vertices within its subject-specific precision mapping-defined spatial extent. We then calculated Pearson correlation coefficients between this mean time series and every cortical vertex across the brain. The resulting correlation maps underwent z-score normalization to standardize the distribution. To allow vertices to be associated with multiple subnetworks based on their functional connectivity strength, each subnetwork was redefined as the set of vertices exceeding a functional-connectivity threshold, which we systematically varied from 0.0 to 2.0. Primary results are reported at a threshold of 1, with additional results provided in the Supplementary Materials. We selected this threshold because, after z-score normalization, a value of 0 corresponds to the mean correlation and 2 represents two standard deviations above the mean— indicative of high connectivity. Higher thresholds retain very few vertices and produce minimal overlap across regions. For each subnetwork pair, overlapping vertices satisfying the threshold criterion for both subnetworks were identified. We calculated the surface area of each cortical vertex to determine both the overlap area and the combined area of the two subnetworks. The overlap ratio, computed as the overlap area divided by the combined area, provided a normalized measure of spatial differentiation.

### Surface area of DMN subnetworks

To quantify the spatial extent of each DMN subnetwork, we first calculated the surface area (in mm^2^) for each cortical vertex. The relative proportion of each network was then calculated by summing the surface areas of its constituent vertices and dividing by the total cortical surface area. After examining age effects on surface area proportion, we further examined developmental changes in the topography of DMN subnetworks. To characterize how the spatial distribution of individualized network parcellations changes with age, we compared individual maps with the Yeo template as a reference, which represents the canonical spatial distribution of each network. By comparing individual network maps against this reference on a vertex-by-vertex basis, we identified vertices that showed different subnetwork assignments across development. For example, by comparing an individual’s DMNC map against the group-level DMNA and DMNB maps, we could determine whether regions that contract from DMNC during development are subsequently occupied by other DMN subnetworks (i.e., DMNA or DMNB) in the reference map.

### Assessing Age-Related Effects on Brain Measures

We employed a general linear model (GLM) to examine the relationship between age and brain measures, including functional segregation, boundary sharpness, spatial overlap and surface area of DMN subnetworks. In these models, age was treated as the independent variable, while the brain measures served as dependent variables. Sex and scanning sites were included as covariates. To account for multiple comparisons, we applied false discovery rate (FDR) correction to the p-values associated with the age variable.

### DMN subnetwork development and cognitive development

We examined whether developmental changes in the segregation and spatial extent of DMN subnetworks are associated with cognitive development. We used scores from the Picture Sequence Memory (PSM) test as a cognitive measure of episodic memory performance. We expected the PSM scores to increase with age, and correlate with the memory-linked DMNC network. In the PSM task, a series of illustrated objects and activities were presented on an iPad in a specific order. The sequence length varied from 6 to 18 pictures depending on the participant’s age. Participants were asked to recall the correct order of the pictures and received points for each correctly recalled adjacent pair. This test assesses the acquisition, storage, and effortful recall of newly learned information. Higher scores indicate better episodic memory performance relative to age-matched peers.

## Acknowledgments

Fonds de recherche du Québec FRQNT 2021-NC-283571 (XJC) Canada First Research Excellence Fund (XJC) Canada Research Chairs program (XJC) Jeanne Timmins Fellowship (YH)

## Author contributions

Conceptualization: YH, JK, XJC Methodology: YH, JK Investigation: YH, JK, HJ, XJC Visualization: YH Supervision: XJC Writing—original draft: YH Writing—review & editing: YH, JK, HJ, XJC

## Competing interests

The authors declare that they have no competing interests.

## Materials & Correspondence

Ying He

## Data and materials availability

Raw data are freely available through the Human Connectome Project, upon completion of a data-usage agreement: [https://nda.nih.gov/general-query.html?q=query=featured-datasets:HCP%20Aging%20and%20Development]. All the analysis and visualization code are available on GitHub: https://github.com/Chai-Neuro-Lab/Segregation-of-DMN-subnetworks

## Notes

### Competing Interest Statement

The authors have declared no competing interest.

