## Supplementary materials for "Development of functional topography of the default mode subnetworks revealed by precision mapping"

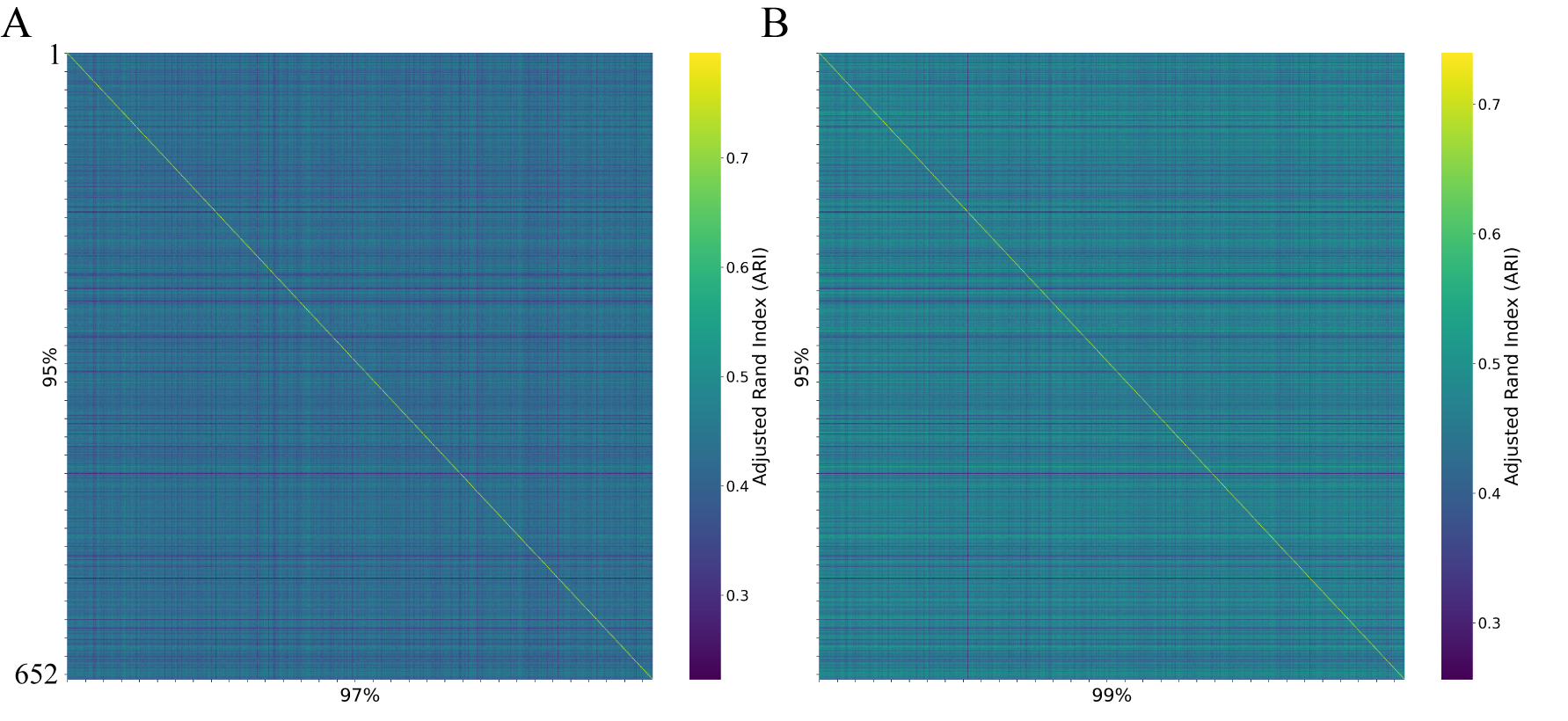

Figure S1. **Threshold Robustness Analysis of Precision Mapping**. A Adjusted Rand Index (ARI) matrix comparing individual precision mappings at 95% versus 97% thresholds. Diagonal elements represent within-subject consistency across thresholds. B. ARI matrix comparing 95% versus 99% threshold mappings with identical diagonal interpretation.

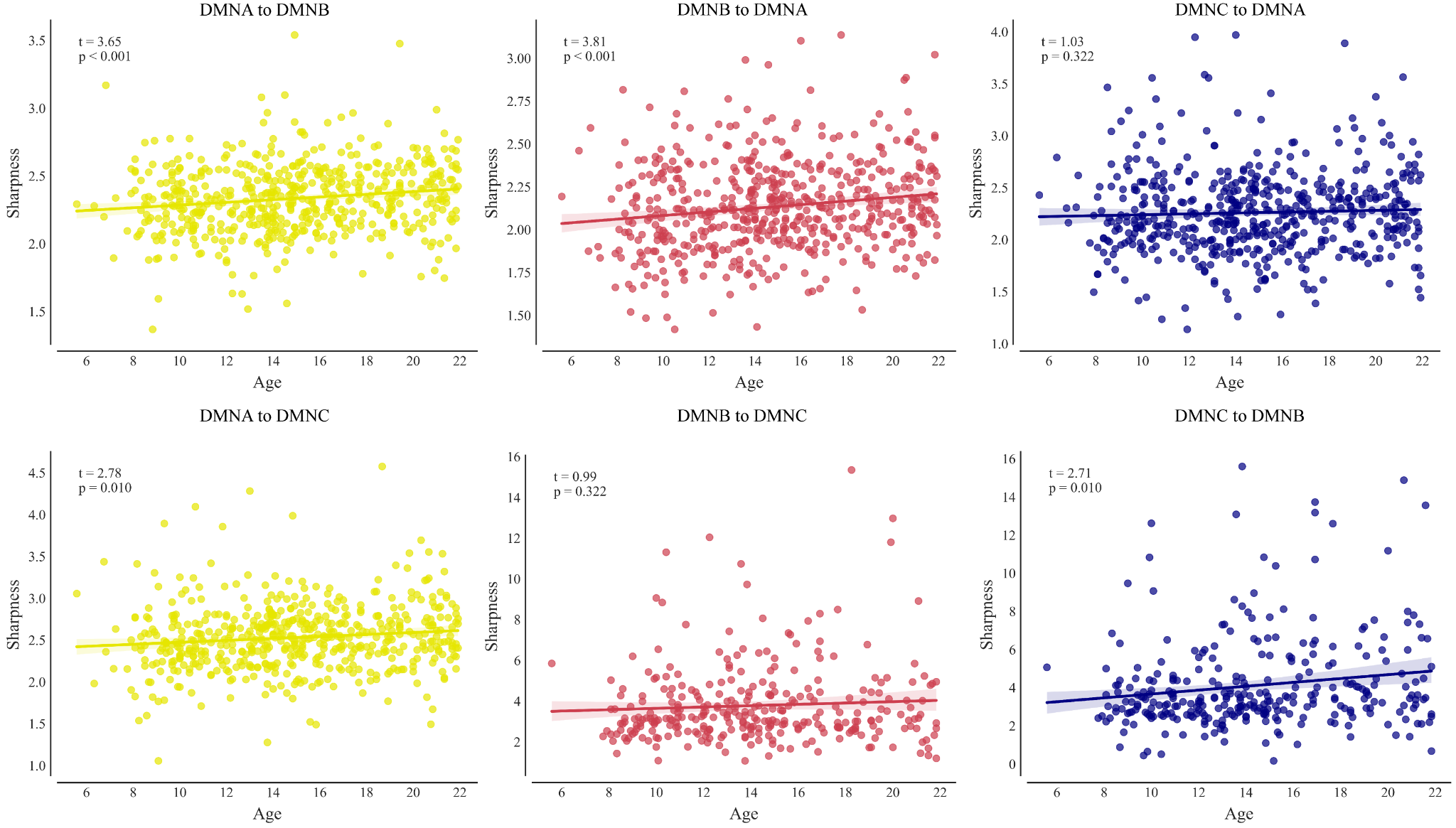

Figure S2. **Age-related changes in boundary sharpness between each pair of DMN subnetworks.** The boundaries between DMNA and both other subnetworks became sharper with age. For DMNB, the boundary with DMNA became sharper, while the boundary with DMNC did not show a significant change. For DMNC, the boundary with DMNB became sharper, whereas the boundary with DMNA remained stable with age.

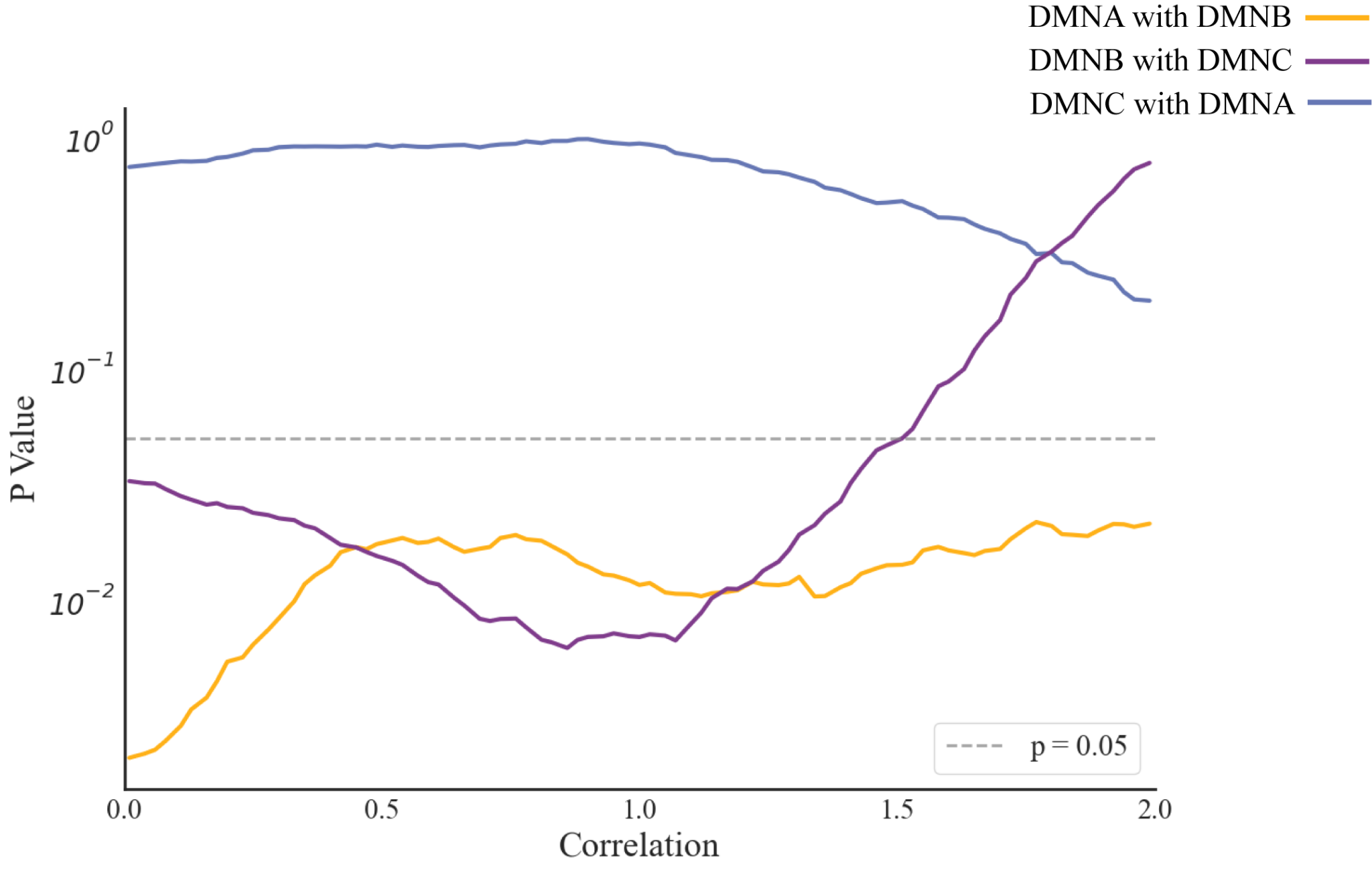

Figure S3. **Statistical Significance of Age-Overlap Surface Area Correlations Across Thresholds.** This figure illustrates the p-values of the correlation between overlap surface area and age across different correlation thresholds (ranging from 0 to 2). Each colored line represents the statistical significance of the relationship for different network overlaps: DMNA with DMNB (orange), DMNC with DMNA (blue), and DMNB with DMNC (purple). The gray dashed line indicates the conventional significance threshold of p = 0.05. The analysis demonstrates that the negative correlation between DMNA and DMNB overlap surface area and age is remarkably robust, maintaining statistical significance across all examined correlation thresholds.

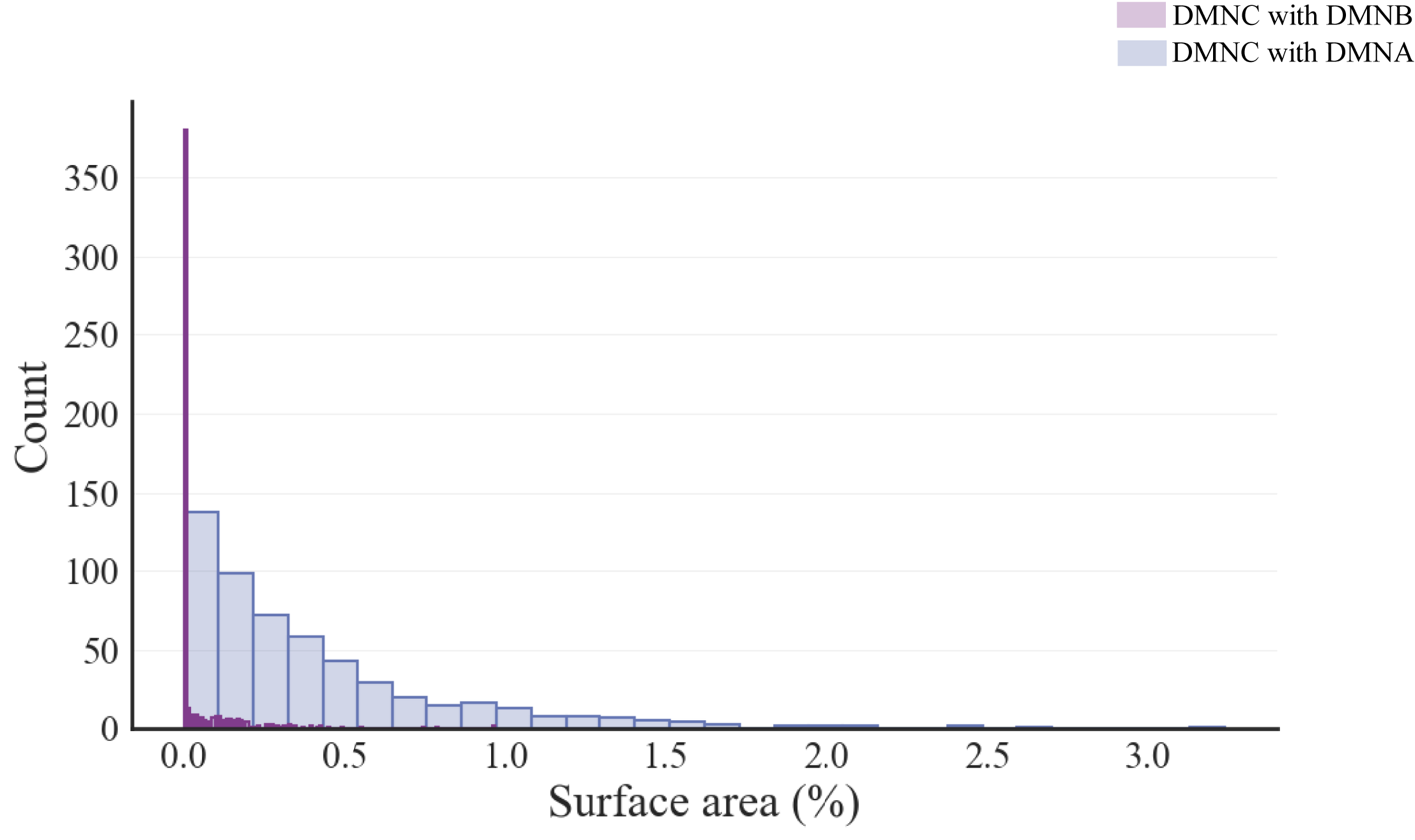

Figure S4. **Reassignment distribution of individual DMNC to group networks.** Histogram quantifying the proportion of individual DMNC surface area reassigned to DMNA and DMNB from template. Y-axis: Subject count exhibiting each reassignment proportion.
